# On the location of a “central retina” in mice

**DOI:** 10.64898/2026.04.16.718979

**Authors:** Alexander Günter, Regine Mühlfriedel, Mathias W. Seeliger

## Abstract

The retinal topography of mammals reflects significant influences of the visual environment. In diurnal species, local specializations, such as the visual streak (VS) for panoramic vision and the *area centralis* or fovea for binocular vision, play a key role in optimizing visual perception and species viability. While the location of these sites is typically considered the retinal center, the definition of a “central retina” is less clear in nocturnal species. In mice, the most frequently used model in ophthalmologic research, the location of a central retina is hardly discernible in retinal images, neither in retinal structure (OCT sections) nor in vascular organization (SLO and angiography). In this study, we compare the murine retina with that of a diurnal rodent, the Mongolian gerbil (MG). We found that the S-opsin transitional zone (OTZ), a region characterized by the change from S-to M-opsin dominance along the dorsoventral opsin gradient in mice, has a similar relative position in the retina to the VS in the Mongolian gerbil, suggesting an evolutionary positional homology between these regions. Further, since the S-opsin-dominant region is optimized for visualizing the sky and the M-opsin-dominant region for visualizing the ground, the OTZ in between –much like the VS– naturally points toward the horizon. We therefore propose considering the OTZ as the position of a “central retinal area” in mice. Determining the anatomical-physiological center is particularly important to obtain meaningful relative measures such as averages across different retinal areas, as the common referencing to the optic nerve head (ONH) in mice does not take into account retinal organization and the eccentric position of the functional center.

## Introduction

Over the course of mammalian evolution, the topographic organization of the retina has adapted to ensure the most effective possible detection of visual stimuli, thereby increasing the survival and reproductive capacity of the species. A key difference lies in the different light environment resulting from nocturnal (night-active) or diurnal (day-active) lifestyles. Nocturnal species usually optimize their light sensitivity under scotopic (dim light) conditions, while diurnal animals typically optimize their visual acuity in relevant areas under photopic (daylight) conditions. A number of factors that further shape these topographies have been described (1, 2). The retinal correlates of visual function are the photoreceptors, rods and cones. Rods possess particularly high light sensitivity and are therefore of paramount importance under scotopic conditions, while cones, in combination with the respective downstream signal processing, enable color vision and high visual acuity under photopic conditions. Important forms of specialized retinal areas such as the macula and fovea, visual streak (VS) and *area centralis* provide increased spatial visual acuity due to a higher cone cell density (3). These are classically considered as central retinal regions in the respective species and are the anatomical basis for the central visual field.

The role of a species as prey or predator is another important factor for the presence of retinal specializations in combination with an appropriate eye position in the head (4). Examples of prey animals include rabbits and rodents such as the Mongolian gerbil (MG), mice, and rats, but also even-toed ungulates such as deer, whose lifestyle requires panoramic vision, which comes at the expense of binocular overlap and depth perception. These requirements favor laterally positioned eyes and a distinct VS as a specialization of the retina. In contrast, the eyes of predators such as cats and dogs are located at the front of the head to enable, in combination with an *area centralis* in addition to the VS, binocular overlap and to improve depth perception. The primate retina enables a very high degree of stereoscopic vision, depth perception and visual acuity, centered on a *fovea centralis* which marks the point of 0° eccentricity in the visual field. (5). Macula and fovea are important reference points for the diagnosis and treatment of many retinal diseases such as diabetic retinopathy and age-related macular degeneration (AMD). (6).

In research on retinal diseases, mice are currently the most popular models, but due to their nocturnal lifestyle, they have a significantly different retinal topography and cone system organization than humans. In particular, they lack a discernible specialization in the ocular fundus, which makes it difficult to determine the location of a “central retina” and the point of 0° eccentricity. Therefore, the centrally located optic nerve head (ONH) is usually used as a reference point instead. However, we found that there are functional clues that can be used to determine a retinal center even when no immediately perceptible structures are present, as is the case with diurnal species. S- and M-opsins are co-expressed in the cones of the mouse along a dorsoventral gradient, with S-opsins being strongly predominant in the ventral retina, thus enhancing the contrast when looking at the sky (presumably for the detection of birds of prey), while M-opsins are predominant in the dorsal retina (Fig. 1A). The region between these two areas, where the dominance of opsin expression shifts, is known as the S-opsin transitional zone (OTZ), which forms the boundary between the upper and lower visual fields (7, 8). While a gradient in cone system organization has also been described in the common shrew, hyenas, guinea pigs and rabbits (9-11), mice additionally coexpress both S- and M-opsins in most cones (Fig. 1A). The remaining cone population consists of “true blue” S-cones, which express only S-opsins and represent an evolutionary ancient pathway in the mammalian retina that may not be image forming (12, 13).

**Figure 1.**
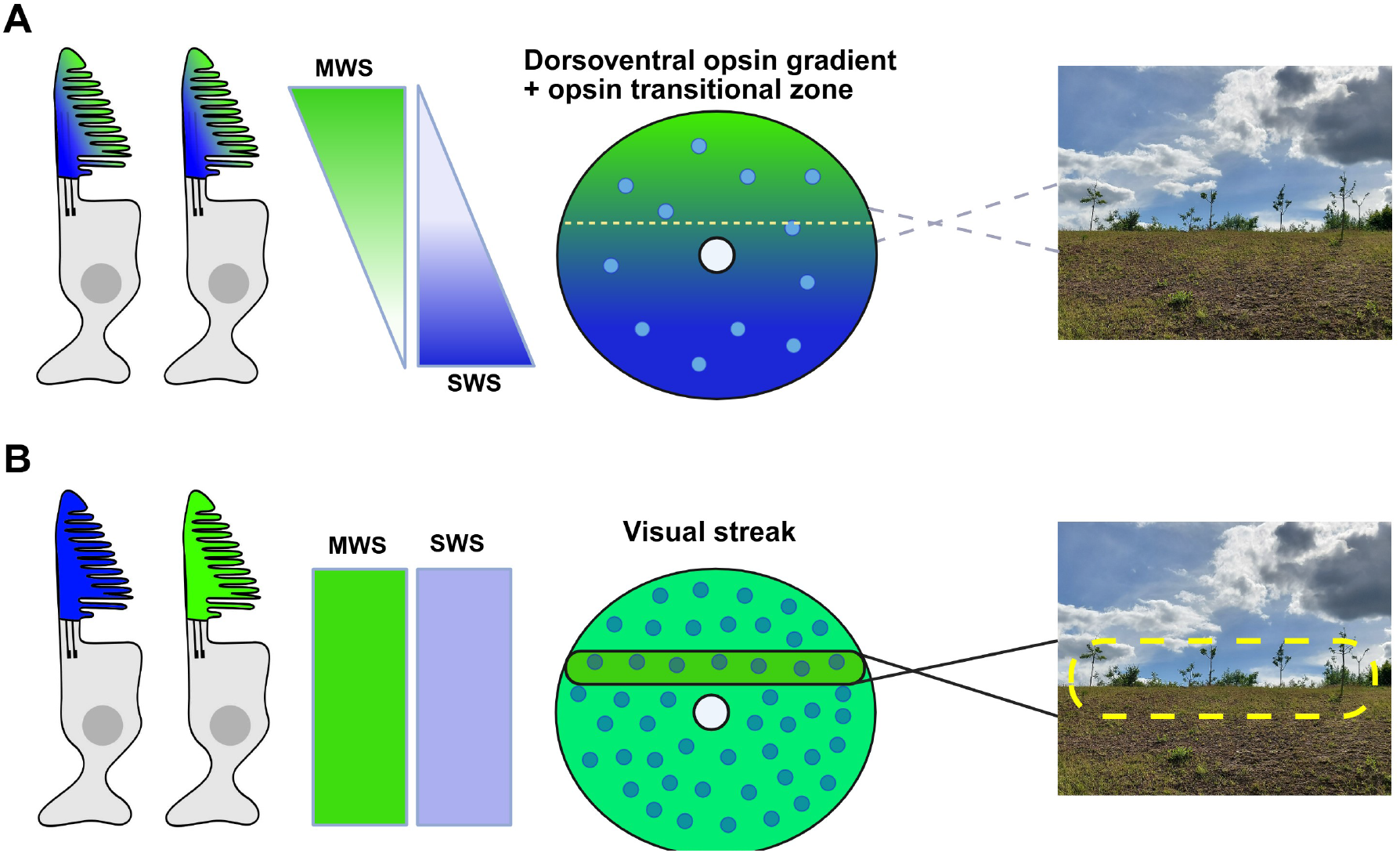
The retinal and cone system organization of **(A)** mice and **(B)** Mongolian gerbils (MGs). **(A)** Mouse cones coexpress M- and S-opsins following a dorsoventral gradient, with a higher fraction of S-opsin in the ventral and of M-opsin in the dorsal retina. A minority of additional “true blue” cones reflect an evolutionarily ancient color vision pathway and express only S-opsin (12). The short wavelength sensitive opsins optimize the ventral retina for the detection of birds of prey due to enhanced contrast against the sky. The S-opsin transitional zone (yellow dotted line) marks the boundary between the upper and lower visual fields. It thus projects towards the horizon similar to a VS, but may not primarily serve as an area of high acuity (grey dotted line). **(B)** MG cones express only one type of opsin per cell, and while the overall distribution of M- and S-opsins is homogeneous throughout the retina, there is a higher density of M-than S-opsins, which is found in most mammalian species (11). The image on the right symbolizes the visual information captured by the VS, i.e., a scan of the horizon. Color bars indicate the distribution of opsins in the retinas of each species, with green representing M-opsins and blue representing S-opsins. In the retinal diagrams, blue circles denote “true blue” S-cones, while the large white circle marks the optic nerve head (ONH) position. Created in BioRender. Mühlfriedel, R. (2026) https://BioRender.com/c3idk4s

To determine whether the OTZ could serve as a potential location for the central retina of the mouse, we compared its location with that of the VS of the MG, a diurnal rodent that is phylogenetically not very distant from mice and likewise belongs to the *muridae* family. MGs are native to the semi-deserts and steppes of Central Asia, whose habitats are characterized by low and sparse vegetation (14). Consequently, they possess a well-developed VS that is ideal for scanning the horizon for predators and locating members of their own group (Fig. 1B). The specialization of its central retinal region is clearly marked in fundus images, both due to a different light reflection/absorption pattern of the VS and due to the organization of the vascular system to avoid crossing vessels impairing visual perception (15). In contrast to mice, the cone system organization in MGs is more comparable to the human situation. For example, M- and S-opsins are homogeneously distributed throughout the retina without evidence of coexpression (Fig. 1B) (15), and the VS features an overall higher density of cones with elongated outer segments relative to the retinal periphery, similar to the border of the macula (16). Also, cone system function is more human-like (15, 16).

In this report, we demonstrate that the location of the anatomical-physiological center in the MG corresponds to the OTZ in mice. We therefore propose using the OTZ in mice as the location of a “central retina” and employing it as a positional reference in research models.

## Materials and Methods

All animal experiments and procedures performed in this study adhere to the ARVO statement for the Use of Animals in ophthalmic and Vision Research and were approved by the competent legal authority (Regierungspräsidium Tübingen, Germany). Mice and MGs were housed in an alternating 12 h light and dark cycle environment with free access to food and water. For the immunohistochemical analysis, 5 adult C57BL/6J mice aged 7-8 postnatal weeks were used. Additionally, we used 5 adult MGs aged 2-3 postnatal months for comparison purposes. Immunohistochemistry, microscopy and image analysis were performed as previously described (15, 16). The following antibodies were used for immunohistochemical assessment: anti-SWS cone opsin (AB5407; 1:300; Merck, Darmstadt, Germany), anti-MWS cone opsin (AB5405; 1:300; Merck), fluorescein isothiocyanate (FITC)-conjugated peanut agglutinin (PNA) (L7381; 1:100; Sigma-Aldrich, St. Louis, MO, USA) and Goat anti-Rabbit, Alexa Fluor 568 (A11036; 1:300; Thermo Fisher Scientific, Karlsruhe, Germany). DAPI was used as nuclear counterstaining. To capture the entire dorsoventral cross-sections, tile imaging was used in conjunction with Z-stack acquisition (Z-planes at 1 μm intervals) using a 10x air objective (N.A. 0.45). The images were subsequently stitched using the ZEN 3.3 (blue edition) software (Carl Zeiss Microscopy), which was also used to measure retinal length and the distances from the optic nerve (ON) to the center of OTZ in mice and VS in MGs.

The limits of the OTZ were determined as follows: (1) The center corresponds to the proximal dorsal retinal region where S-opsin is greatly reduced and no longer detectable in nearby cones. Quantitatively, the center of the OTZ was determined by the first S-opsin positive cone whose nearest neighboring cone of the same type lies at a distance of ≥ 25 µm in the cross-section. This position was then used as a reference point for analyzing distances from the ON. (2) The initial limit of the OTZ was determined by the location where M-opsin begin to increase in expression in cones and S-opsin density is not as prevalent as in the ventral counterpart, which is ∼200 µm dorsal from the ON. (3) The final limit of the OTZ was determined by the last S-opsin positive cone whose nearest cone of the same type lies at a distance of ≥ 50 µm in the cross-section. In MG, the center of the VS was defined as the midpoint of the region highlighted in Figure 2B, characterized by stronger PNA staining. The dorsoventral retinal length was measured along the retinal curvature in cross-sections. For histological analysis, only sections with an apparent ON were selected. For the statistical analysis of immunohistochemical data, we used two-tailed unpaired t-tests for two groups.

**Figure 2.**
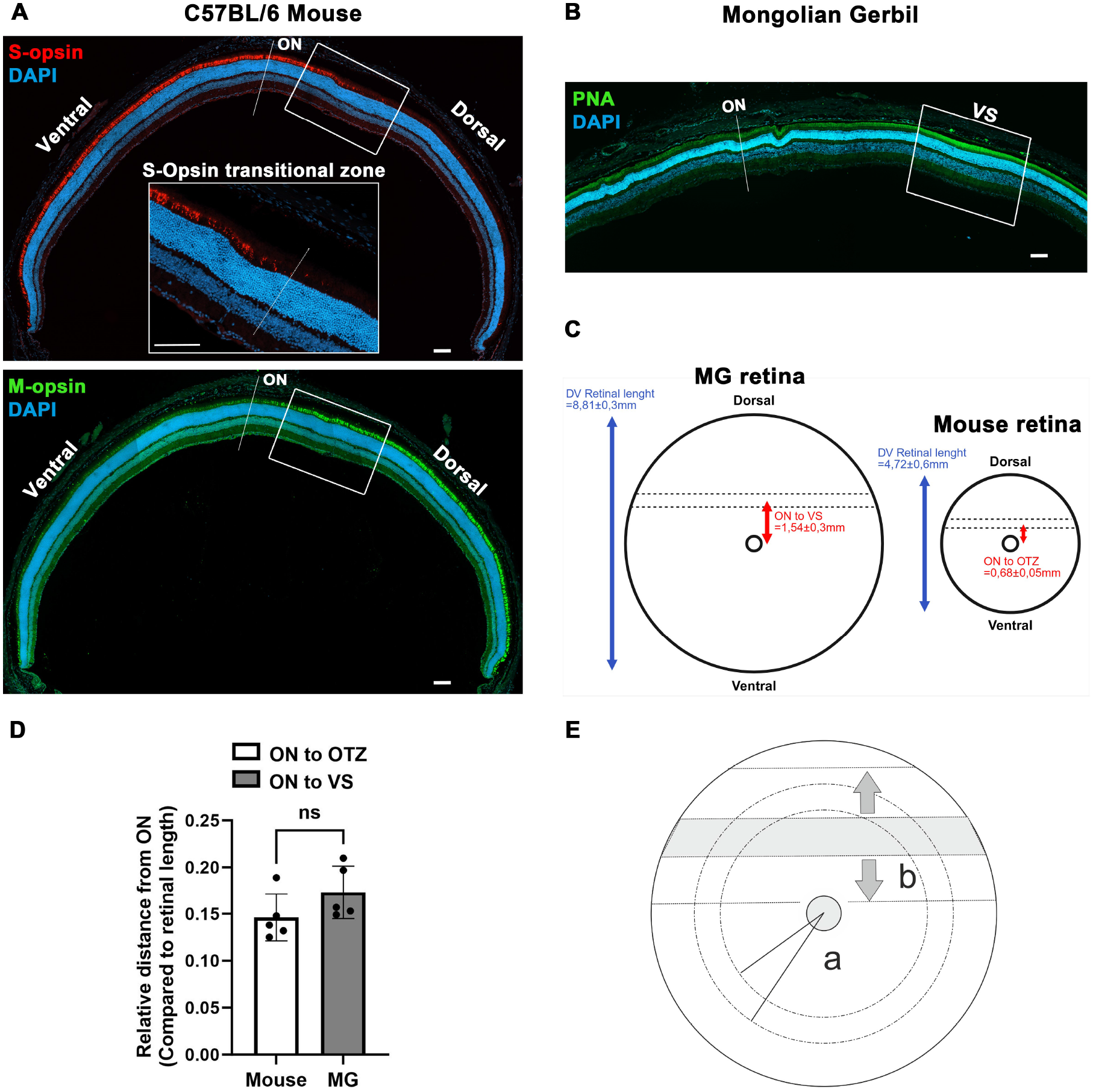
The S-opsin transitional zone and the VS. **(A)** Dorsoventral section of the C57BL/6 mouse retina showing the distribution of S- and M-opsins. S-opsins are highly expressed in the ventral retina with only a few cones showing a positive staining in the dorsal portion, which correspond to the “true blue” cone subtype. In contrast, M-opsins are predominantly expressed in the dorsal retina and show reduced expression ventrally, likely due to the high level of coexpression with S-opsins in this region. Boxes highlight the OTZ region, which is characterized by an abrupt reduction in S-opsin density in the proximal dorsal retina. The inset shows an enlargement of the OTZ; scale bar = 100 µm **(B)** PNA highlights the cone OSs in the MG retina. Expression is strongest in the VS, reflecting the higher cone density and the elongated OSs in this region and its suitability as a VS marker. ON = optic nerve; VS = Visual streak; scale bar = 100 µm. **(C)** Schematic representation of the distances from the ON to the OTZ or the VS in relation to the total dorsoventral retinal length. **(D)** Analysis of the relative distance from the ON to each specialized retinal region (OTZ in mice and VS in MG), expressed as a proportion of the total dorsoventral retinal length (n = 5 eyes). There are no significant differences in the relative distances between both species. Error bars represent the standard deviation; n.s. = *p* > 0.05. **(E)** (a) Concentric ring areas based on the ONH as reference point include physiologically very diverse photoreceptor populations and should therefore not be used for averages across retinal cells. (b) The OTZ in mice forms an eccentric reference line similar to the VS in MGs as the central position in the retina. Arrows indicate dorsal and ventral positions with the same relative eccentricity.

## Results

To determine the position of the OTZ in the opsin-coexpressing cones, we performed immunohistochemical stainings of S- and M-opsins in the retina of C57BL/6 mice. In the ventral and in the proximal region of the dorsal retina, we found a strong staining of S-opsins (Fig. 2A, top). In the dorsal retina beyond this region, only a few S-opsin-positive cones were observed, representing the class of “true blue” cones. Conversely, M-opsins were dominant in the dorsal retina, but were also present to a lesser degree in the ventral retina (Fig. 2A, bottom). The OTZ is highlighted as the region where the S-opsin gradient ceases and the M-opsin gradient rises in the proximal dorsal retina (Boxes and Inset in Fig. 2A). The center of the OTZ may be determined as the position where nearby cones no longer stain for S-opsin (Inset in Fig. 2A, dotted line).

To assess whether the position of the OTZ corresponds to the location of the anatomical-physiological center in the MG, we compared the relative distances from the ON to the OTZ in mice and to the VS in MGs, expressed as a proportion of the total dorsoventral retinal length (Fig. 2B). The average distance from the ON to the VS in MGs was 1.54±0.3mm and the distance from ON to OTZ in mice was 0.68±0.05mm, while the average retinal length was 8.81±0.3 mm in MGs and 4.72±0.6 mm in mice (Fig. 2C). These measurements did not reveal any significant difference between the relative ON-OTZ distance (mice, 0.14±0.02) and the ON-VS distance (MG, 0.17±0.03) (Fig. 2D). In other words, the eccentricity of the OTZ (the distance between ON and OTZ center) is approximately 14% of the total retinal length in mice, compared to 17% for the eccentricity of the VS (the distance between ON and VS center) in MGs. These data suggest a positional homology between the OTZ and the VS, which indicates that both structures either originated from a common ancestor or developed during evolution due to functional necessities. Consequently, as the location of the OTZ in mice corresponds well to that of the VS in MGs, we suggest its use as reference for the retinal center. Referencing the correct anatomical-physiological center is particularly important to obtain meaningful relative measurements related to the cone system, as the common referencing to the ONH in mice does not take into account retinal organization and the eccentric position of the physiological center, as highlighted in Fig 2E.

## Discussion

Cone cells existed before rods if we look into a period of more than 500 million years ago. A jawless proto-vertebrate presented cone-like photoreceptors expressing SWS or LWS opsins. Two rounds of whole genome duplications resulted in the formation of four cone-opsin classes and one rod class specialized for low intensities (17). Until the period of 150-200 million years ago, early mammals were believed to live with a predominant diurnal lifestyle. This changed with the domination of archosaurs, that forced them to change to nocturnality to avoid becoming prey (18). During this time, early mammals lost two classes of cone opsins and underwent a reduction in the total number of cone cells. The extinction of dinosaurs reopened the option of a diurnal lifestyle and stimulated the development of different types of retinal organizations. The Rodentia order is a good example in terms of the variability of retinal organization including currently diurnal, nocturnal, and crepuscular lifestyles (19).

Here, we took advantage of the different retinal organizations in two rodents with different lifestyles, the nocturnal house mouse (*mus musculus*) and the diurnal Mongolian gerbil (*Meriones unguiculatus*), to determine the position of the anatomical-physiological center of the retina in the former. We found that the OTZ in the mouse occupies the same relative position in the retina as the VS in MGs, suggesting that the OTZ may represent a positional homology to the VS (20).

The mouse cone system has traditionally been considered relatively obsolete for visual function, but this perspective has changed in recent years. The dorsoventral gradient distribution of opsins in the mouse retina is motivated by their role as prey in nature. S-opsins in the ventral retina were shown to be biased towards dark stimuli, which converges with the prevalence of dark and achromatic contrasts in the sky. This is a key feature that assists mice in the early detection of incoming aerial predators (21). In terms of chromatic processing, the OTZ presents unique properties. According to the immunohistochemical staining of the mouse retina presented in Fig. 2A., this is a region where S- and M-opsin densities rapidly change over a short distance, without clear dominance of either opsin type, an organization not observed in any other retinal region. Further, the OTZ was shown to present a higher abundance of color-opponent retinal ganglion cells (8). Additionally, the OTZ location coincides with the horizontal meridian from mice, as revealed by S-opsin staining, indicating that this region marks the boundary between the upper and lower visual fields (7).

In the light of these facts and based on our data, we propose that the OTZ is considered the reference for the central position in the mouse retina. While it is widely accepted that the fovea is the center of the human retina (6), standardized reference points in nocturnal rodents are rare. Often, the ONH that marks the geographical center of whole-mounts and *in vivo* images, is used in the absence of a meaningful, recognizable alternative. However, this not only reduces the comparability of the results of disease models with those of human patients, but also leads to problems in averaging over equidistant regions in the analysis. Therefore, an agreement on an anatomically and physiologically correct retinal center in rodents such as the OTZ is highly desirable.

## Author contributions

Alexander Günter: Conceptualization, data curation, formal analysis, investigation, validation, methodology, visualization, writing – original draft preparation. Regine Mühlfriedel: Funding acquisition, project administration, resources. Mathias Seeliger: Conceptualization, funding acquisition, project administration, resources, supervision, visualization, writing – review & editing.

## Funding

This research was funded by the German Ministry for Education and Research (BMBF; TargetRD 16GW02678) and the German Research Council (DFG; SE 837/12-1). We acknowledge the support from the Open Access Publication Fund of the University of Tübingen.

## Competing interest

The authors have declared that no competing interests exist.

## Acknowledgements

We thank Gudrun Utz for the technical assistance and Dr. Yu Zhu for assessing the replicability of data analysis.

## Notes

### Competing Interest Statement

The authors have declared no competing interest.

